# Roost: Avian Influenza Host Genomics Resource

**DOI:** 10.64898/2026.11.01.763514

**Authors:** Joseph Guhlin, Peter K. Dearden

## Abstract

Highly pathogenic avian influenza (HPAI) causes mass mortality in poultry and wildlife, and puts already threatened species at further risk. Understanding differences in host susceptibility requires evidence currently scattered across publications, surveillance reports and genomic resources. We present Roost, a web resource that brings together more than 1,300 publications and reports, surveillance detections, and a catalogue of over 2,000 avian genome assemblies. Large language model-assisted literature analysis and comparative genomics link species, genes and molecular mechanisms to their supporting evidence. Around 51% of species with HPAI detections are not represented in the analysed corpus. Research concentrates on poultry, model organisms, and a small set of immune pathways. Research tracks neither mortality magnitude nor extinction risk; species with the largest recorded die-offs and threatened hosts alike remain largely unstudied. Roost makes this evidence, along with genome analyses, accessible to researchers and conservation practitioners, connecting documented disease with host biology and highlighting priorities for sequencing and functional research. Roost is available at http://josephguhlin.com/Roost

## Main

Highly Pathogenic Avian Influenza (HPAI) represents a serious worldwide threat to agriculture, wildlife and potentially to human health. With at least 718 species affected and over 44,871 detection records spanning more than 210 million birds. Likely undercounted in wild species, this threat is widespread, damaging, and increasing. 73 threatened avian species are among detected HPAI hosts^1^, risking extinction of fragile populations. Because host species vary in susceptibility, disease severity, and transmission risk, pathogen surveillance alone cannot predict disease emergence and its consequences.

Genome assembly has become routine^2–5^. Research on HPAI, including disease studies, functional studies, gene annotations, and mechanistic findings, continues to emerge rapidly. With frequent new detection events and new HPAI host species being added, the field is a dynamic, changing landscape.

Pathogen genomic resources and surveillance data are mature, including viral sequence data-bases, pathogen phylogenetic resources, surveillance dashboards, strain tracking and more^6–13^. Host genetic data is scattered across detection reports and individual studies. Researchers have no easy way to query the H5N1 literature about specific genes, risk factors, host biology and genomics, and species-specific resources. Likewise, conservation managers have no easy way to identify a species’ risk factors unless it has been directly studied.

In recent years, major efforts have sequenced the genomes of multiple species across the tree of life^14–16^. With over 2,000 non-redundant genome assemblies across Aves, this resource lets us predict risk and susceptibility based on phylogenetic information and the presence or absence of key genes. While only a prediction, it can help prioritize resources, such as vaccination or quarantine, more effectively and highlight where additional research should focus for higher impact in addressing this panzootic threat.

We present Roost, a resource for the host genetics side of HPAI across all Aves integrating over 1,300 publications, genomic analyses of over 2,000 nonredundant genomes, and species-level information for 718 species. Our goals are to synthesise host genetic and molecular evidence, connect literature-derived biology with comparative avian genomics, and quantify where host-genetic knowledge is concentrated and where major research gaps remain. Roost serves as a central repository for HPAI information with a non-exclusive focus on host genetics in Aves.

## Results

### Roost is continuously updated; numbers reflect those at time of submission, live website may differ

We analyzed 1,366 papers, 44,871 viral detections covering 718 species across 200 million individual animals from WOAH WAHIS^11^, USGS WHISPers^17^, FAO *EMPRES*-i^18^, SA OCIMS^19^, New Zealand MPI / Biosecurity NZ^20^, Wildlife Health Australia^21^, 2,093 genomes and 302 gene annotations from NCBI^22,23^, additional protein sequences from UniProt^24^ and ImmunomeDB^25^, taxonomy from Birds of the World, Clements Taxonomy^26^, conservation status from IUCN Red List^1^, additional taxonomic name resolution from GBIF^27^, protein 3D structures from *RCSB PDB*^28^, and additional facts, citations, and knowledge from the book Avian Immunology^29^ (see Figure 1.a).

**Figure 1:**
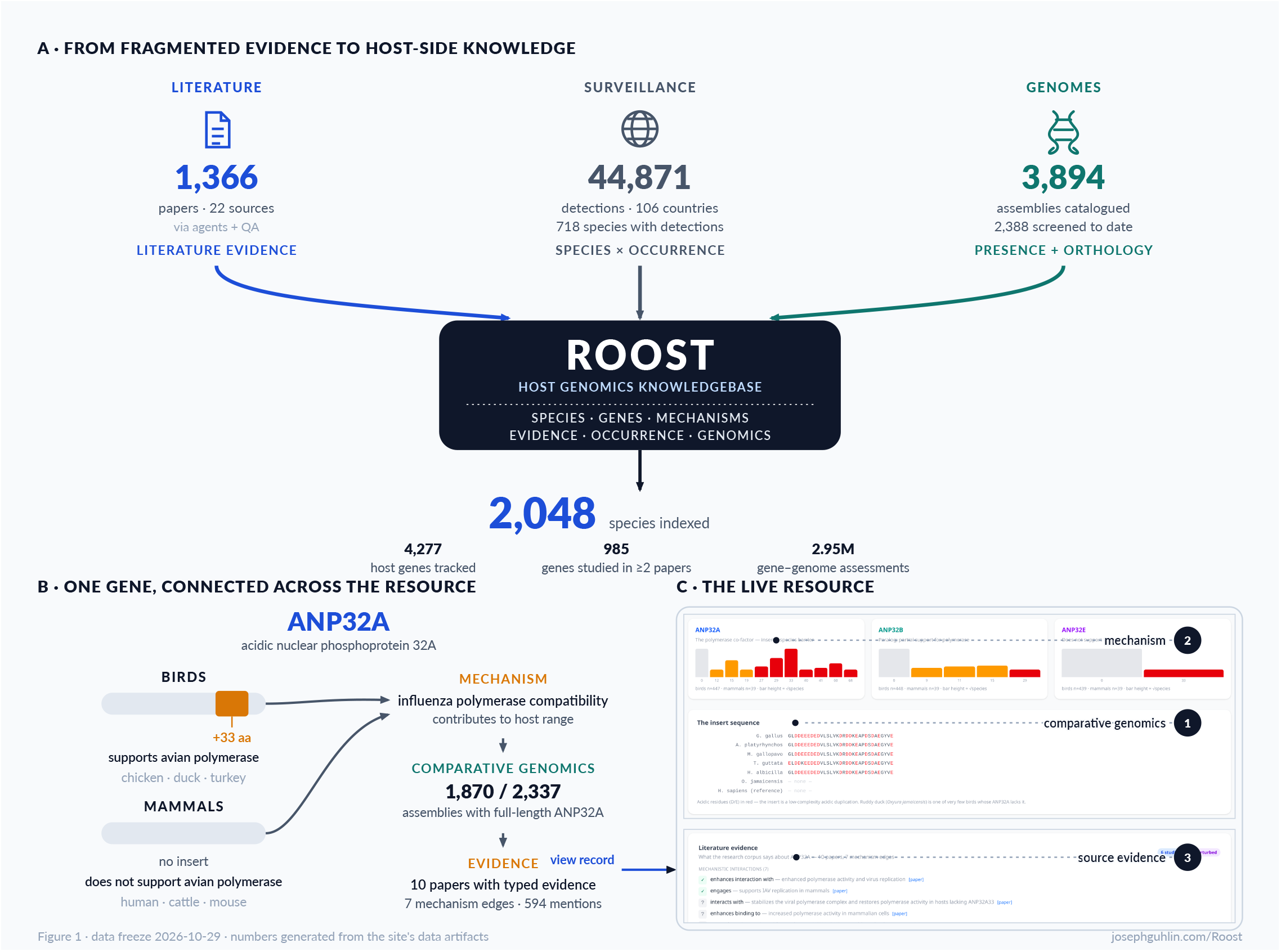
A. Roost is generated from multiple data sources; most importantly, primary scientific literature, detection events, and genomes. B. Cladebarrier gene *ANP32A* has a 33-amino-acid insertion that mammals lack, which previously limited the range of hosts affected by HPAI. Roost identifies this mechanism and the presence of this gene in assayed genomes, with links to primary literature for evidence. C. For a gene-level view, you can see multiple sequence alignments, the presence or absence of that gene in other genomes, and the evidential statements with links to the original papers.

### Integrating host genetics, molecular mechanisms, and comparative ge nomics

Literature analyses identified 1,519 host genes, 3,606 functional relationships, and 721 or-thogroups containing all 992 host panel genes, across 1,366 citations, including 1,152 with typed gene claims and 519 with mechanisms. We detect, on average, 83.5% (median 83%, *IQR* 78– 89%) of Tier 1 host dependencies (e.g. *ANP32A/B/E*; Supplemental Table 1) and 87.6% (median 89%) of total host genes across 2,388 genomes.

Literature is processed and parsed using a locally hosted large language model (LLM). The literature, including published scientific articles, preprints, theses, news reports, and government reports, is categorised; species and referenced genes are extracted; primary findings are extracted iteratively; and relevant databases are updated. Provenance is kept at each step, and multiple iterations are performed to reduce hallucinations. The LLM organises, synthesises, and summarises evidence.

As an example of the summarized LLM evidence, the gene page for *ANP32A* states, “*ANP32A* is an acidic nuclear phosphoprotein that acts as an essential host cofactor for influenza A virus RNA-dependent RNA polymerase. In the context of H5N1 avian influenza, *ANP32A* binds the viral *PB2* subunit and supports vRNP assembly, transcription and genome replication in avian cells. Adaptation of H5N1 to mammals and to cattle in particular involves altered polymerase-*ANP32A* interactions, with bovine *ANP32A* becoming the primary cofactor for adapted viruses.” This statement is generated from 41 sources. Research confirms mammalian adaptation requires altered *ANP32A* interactions;^30^ and *ANP32A* are primarily the host barriers^31^ (Figure 1.b). Meanwhile, gene-edited chickens with altered amino acids in ANP32A do show some resistance^32^.

### Literature-wide synthesis identifies the molecular landscapes of host re sponse

We have identified 4,277 distinct host genes implicated across 1,366 papers, plus 617 non-gene entries in the registry, such as conceptual categories, miRNAs, *MHC* alleles, viral subtypes, and mutations. Papers are linked to 3,502 genes through extracted study roles. Singletons, in which a gene appears in only one paper, account for 71.9% (2,517) of genes and are generally excluded from our gene panel. 17.3% of genes are linked to 2 to 3 papers, and 1.4% (50) genes are supported by ≥20 papers. The remaining genes are panel-only, with no supporting literature (297), but are included due to their relevance as inferred from the UniProt/GO immune backbone.

Across 3,502 studied host genes, the ten most-studied — *IFNB1* (102 papers), *TNF* (100), *IFNG* (95), *IL6* (94), *MX1* (93), *DDX58* (83), *IL1B* (69), *IRF7* (61), *GAPDH* (51) and *ISG15* (50) — the interferon core and cytokine response — account for 9.5% of all gene-paper links, and 80% of the literature is spread across 1,827 genes, with a median of one paper per gene (Figure 2.a,b). Fifty genes have twenty or more studies, spanning interferon signalling, *RIG-I*/*MDA5* sensing, and the *OAS-RNaseL* axis. Most of our understanding of genes affected by HPAI comes from single studies of individual genes and often describes a hypothetical interaction. In these single studies, evidence classes are crucial: singleton genes carry perturbation evidence, where a genetic experiment tests the predicted host-gene-virus interaction only 3% of the time, versus 76% for genes with twenty or more studies.

**Figure 2:**
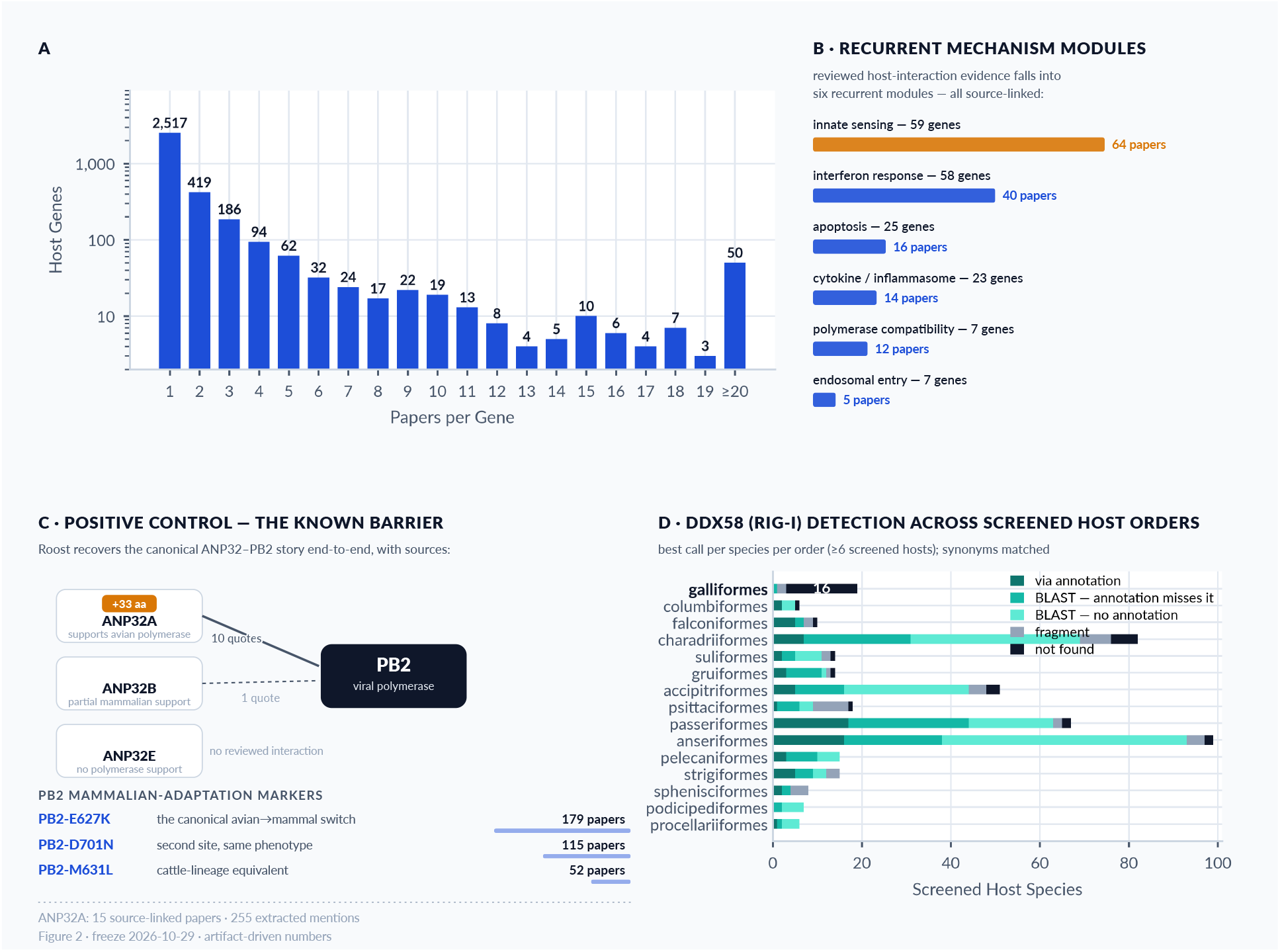
A. Most genes mentioned in the assayed literature appear in only a single paper (2,517 of 3,502 study-subject genes, 71.9%); only 985 appear in two or more, while the most-studied genes (*IFNB1, TNF, IFNG, IL6, MX1, DDX58*) each rest on 83–102 papers. B. The 299 reviewed mechanism edges classify into six recurrent modules (261 edges in-module); innate sensing is the deepest (64 papers, 102 edges). C. *ANP32A* is the studied species barrier, and Roost recovers the canonical *ANP32A*–*PB2* interaction end-to-end: the mammalian-adaptation markers *PB2*-*E627K* (179 papers), *D701N* (115) and the cattle-lineage *M631L* (52) are each recoverable with paper-level provenance. D. *RIG-I* (*DDX58*) is a known sensing gene whose loss has been extensively studied; high lethality results from its absence in chicken. This loss is reflected across most galliformes. This well-studied gene is present in most clades.

### Comparative genomics extends host biology across avian diversity

Approximately 19% of bird species have genomes, with 3% having chromosomal-level assemblies (18% of available genomes). Available avian genomes are dominated by Passeriformes (784, 37%), followed by parrots (217), raptors (161), shorebirds (151), and waterfowl (144). The intersection of those also affected by HPAI skews the other way, with waterfowl (Anseriformes, 100) as primary, followed by shorebirds and gulls (Charadriiformes, 83), raptors (51), and passerines (65). These encompass the reservoir (waterfowl), coastal vectors (primarily Charadriiformes) and raptors (scavenging spillover). While the waterfowl reservoir is nearly covered for genomes (100 of 111 detected host species), the same is not true further down the transmission chain. Of 208 detected host species with no genome assembly in our queried repositories, the gaps are also concentrated: 47 shorebirds and gulls, 44 colonial seabirds (pelicans, cormorants, gannets and albatrosses), and 51 passerines. These species, which have the largest mortality events, such as the Cape cormorant (26,540 dead^33^) and California gull (20,055^17^), are least sequenced.

We extracted genomes and gene annotations from *NCBI*^22,23^. Only 302 assemblies (8%) covering 255 species (12%) have predicted gene annotations. We searched our gene panel across all genomes for both predicted genes, when available, and the genome. To maximize coverage, we used no quality threshold to filter or remove individuals, but we did not analyze redundant or prior genome versions. This connects gene panel entries to predicted genes and genomes and provides presence/absence status for each gene panel entry in each sequenced species.

For example, you can track *ANP32A*, the host barrier gene, across species, including sequence alignment and literature evidence (see Figure 1.c Figure 2.c). This extends to phylogeny, and the phylogeny viewer shows each gene’s status across species and clades, enabling broad pattern detection to infer status in unsequenced species. This is seen in Figure 2.d for *RIG*-*I*, a well-studied gene primarily absent in galliformes, but present in most clades.

### Host-disease research is limited

Genomic resources are no longer a limitation to discovery; while additional sequencing of at-risk species is still needed, future research effort should focus on infectious mechanism research. HPAI research is intensely concentrated on a small number of species: the ten most-studied species — Chicken (369 papers), Human (224), Mouse (166), Duck (generic, 112), Cattle (94), Turkey (87), Goose (77), Ferret (60), Mallard (60), Red Fox (39) account for 34% of all species-paper links, and 80% of the literature concerns just 181 species, with most species appearing in just a single paper (Figure 3.a). The remaining research effort is split between reservoir species (Anseriformes, 482 papers), poultry (Galliformes, 495), coastal vectors (Charadriiformes, 309), raptors (142) and passerines (50). Most worryingly, 366 of the 718 detected HPAI host species (51%) have never been studied in our corpus, and some of the most common HPAI hosts are largely unstudied (Figure 3.b).

**Figure 3:**
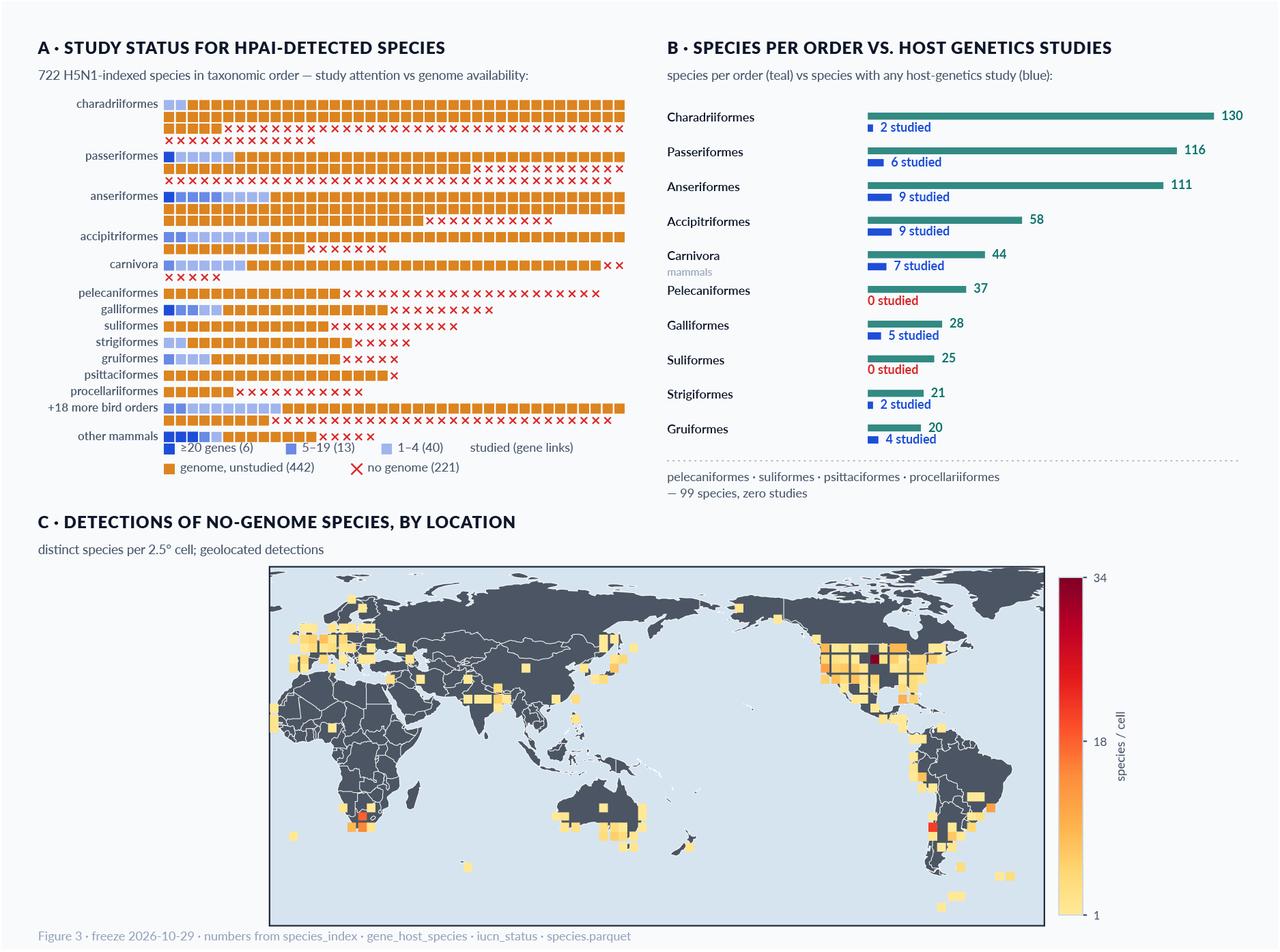
A. Study attention for species with known HPAI detections. Boxes indicate an available genome, while blue indicates the number of genes studied in that species. Orange indicates species with an available genome but no host-gene research related to *HPAI*. B. Species per order (teal) against host-genetics study attention (blue): 4 of the larger bird orders have zero studied species (Pelecaniformes, Suliformes, Psittaciformes, Procellariiformes), and within the studied set, 40 of 59 studied species are named by ≤4 genes, while only 6 species carry ≥20 gene links. C. Geographic distribution of the no-genome gap: every H5N1 detection of a no-genome species with coordinates (1,239 records) binned into 2.5° cells, coloured by distinct species per cell (Pacific-centred). The gap concentrates along coasts and wetland flyways: hottest cells are the US Great Plains (34 species), South Africa’s Western Cape (21), Patagonia (18) and south-eastern Brazil (12); 62 countries carry at least one such species. All numbers are generated from the same artifacts as the live site (data freeze 2026-10-29).

Of the 718 species detected with HPAI, 68 are well-studied (≥3 corpus papers) and have genome resources (Figure 3.a). These are mainly poultry and waterfowl. A further thirteen species are biologically well-studied yet lack a genome assembly in our queried set: kelp gull, Peruvian pelican, Cape cormorant, black-necked swan, South American tern and marine otter should be sequencing priorities. The remaining 208 detected species are neither studied nor sequenced, a gap dominated by gulls and terns, which include species with the largest mortality events, including the California gull and the swift tern.

Research papers are concentrated in a small area of the phylogeny of birds, from 13.8 papers per detected host species in Galliformes, through waterfowl (3.1) and coastal Charadriiformes (1.8), down to raptors (1.5), pelicans and allies (1.1), and passerines at just 0.4 papers per host species despite 116 detected species. Entire families are effectively unresearched: among parrots (Psittacidae), thrushes (Turdidae) and plovers (Charadriidae), only two or three species of sixteen, fifteen and thirteen detected hosts respectively have ever been the subject of a paper providing information that may help our HPAI response, and ibises (Threskiornithidae) only marginally better at four of twelve. Genomic representation partly compensates where sequencing programs have independently targeted biodiversity, waterfowl (100 of 111 hosts with assemblies) and raptors (51 of 58) chief among them, but procellariid seabirds, with five genomes for fifteen detected hosts, remain doubly underrepresented in our research efforts. Research attention, unsurprisingly, tracks agricultural and model-system value rather than either exposure or mortality, a bias that is invisible at the species level and only becomes actionable when mapped onto the phylogeny.

The gap between documented disease and research is not a data problem. Among the 576 detected host species with fewer than three research papers are ten whose combined mortality exceeds 517,000 counted dead birds, led by Ross’s goose (91,450), the white-fronted goose (78,410) and the American white pelican (76,050), each with surveillance records numbering in the dozens to hundreds. Surveillance has documented these die-offs in detail; research has simply never followed. Nor does conservation status steer attention: the share of effectively unstudied hosts is nearly flat across *IUCN* categories — 56 percent of threatened host species (41 of 73) against 51 percent of Least Concern species — with the largest mortality events falling almost entirely on Least Concern taxa. Research effort in HPAI host biology tracks neither mortality magnitude nor extinction risk, leaving both the panzootic’s biggest victims and its most vulnerable species equally unstudied.

## Discussion

Despite high-quality resources for HPAI pathogen surveillance, HPAI lacks a host genetics counterpart. HPAI is unique because it poses a potential risk not only to human health but also to agriculture and wildlife, and continues to adapt to new species^34–36^. Roost fills that gap, bringing together avian genomic resources and HPAI literature to support further research. Previously, ImmunomeDB^25^ was the central source for avian immune genes. However, it focused on pathogens and immunology rather than host genetics, and it was a general-purpose resource rather than one focused on a single disease outbreak. *GISAID*^6^, Nextstrain^7^, and other surveillance dashboards are pathogen-centric. No resource currently integrates host genetics, detections, and literature.

Roost identifies species with confirmed HPAI cases, aggregates detections from multiple public sources, and synthesizes evidence, leveraging decades of genomics and immunology research to produce a centralized data source for researchers and wildlife managers. Comparative genomics makes it possible to translate knowledge between species.^35,37–41^

For literature extraction, we use a multi-pass LLM approach: each paper is registered and screened, a primary pass extracts genes, species, mutations, quantified findings, and quotegrounded mechanisms with verbatim provenance, and an independent adjudication pass re-reads every item against its quote — currently 86% pass as faithful, and failures are flagged in the released data rather than silently dropped. Deterministic checks reconcile 21 distinct data-source totals with the website totals on each rebuild, surfacing uncertain records for human review. All analysis models were hosted locally, preventing copyrighted literature from being used for training without authors’ consent, on servers in Aotearoa New Zealand that draw primarily on renewable energy. Together, this shows how literature synthesis at corpus scale can be performed ethically and responsibly, with every claim traceable and every failure marked.

To reduce potential hallucinations, we queried the LLM multiple times with multiple prompts, varying the granularity of information and the questions, and asked for structured outputs (i.e., does this paper support or reject this statement; respond only with a numeric score). Regardless, LLM extraction may still contain errors, and Roost’s strength is that nearly all statements can be traced back to the original statements and their original publications. Roost-generated knowledge should be considered auditable, not a primary source. Assembly quality affects the gene-presence matrix. This is reflected in missing genes, which could be genuinely missing, unassembled, unsequenced, or affected by assembly errors. Literature statistics are sensitive to how names are joined: species-level paper counts move slightly between corpus versions as names are canonicalised and new papers arrive; “duck”, for instance, is the corpus’s generic term rather than specifically *A. platyrhynchos*.

Against the global threat of HPAI (Figure 3.c), multiple technologies and angles have risen to understand this disease. Genomic technologies, expression analyses, forward and reverse genetics, agricultural and wildlife surveys, researchers from multiple fields and backgrounds, and more have all contributed towards confronting this problem. LLMs help us digest and process data at previously impossible scales, and Roost allows us to layer it in innovative ways.

For conservation practitioners, Roost provides additional ways to gauge disease risk. For unsequenced species, the gene panel presence matrix alongside the phylogenetic viewer for gene presence/absence can help inform decisions and estimate susceptibility. Several species are highvalue targets for genomic sequencing: kelp gull, Peruvian pelican, Cape cormorant, black-necked swan, South American tern, and marine otter would connect field evidence to a genome, and ultimately the genes those species contain. Further genomic work should prioritize high-quality gene annotations for avian species regardless of HPAI detection, and sequence species with positive detections but no genome sequence. Roost reveals these targets for future genomic sequencing, but also contains information useful to conservation practitioners and wildlife rangers, diseasespread modelers, and those doing broad phylogenetic analysis of HPAI, which did not exist in an accessible format before but was spread across scientific literature and detection events reported in the media and by government agencies.

## Conclusions

Roost brings decades of genomics, immunology and disease surveillance together to support research on avian influenza. The assembled evidence exposes a substantial gap between documented disease and our understanding of host biology, particularly in wild species. Available genomes create opportunities to extend research across avian diversity, while gaps in sequencing and functional evidence highlight where further work is needed. As HPAI continues to threaten agriculture and wildlife, Roost provides an accessible, regularly updated resource that connects new findings with existing knowledge and helps researchers and conservation practitioners prioritise their efforts.

## Methods

Detailed, up-to-date methods are available at http://josephguhlin.com/Roost/#/methods.

### Website

Roost is served as a Svelte^42^ single-page web application. All data ships as versioned static artefacts (*JSON*, parquet and DuckDB^43^ files served over *HTTP* and queried client-side via DuckDB-WASM), so the site needs no server-side database and every displayed number traces to a generated artefact. A deterministic cross-check build (“by the numbers”) reconciles 21 artifactvs-page totals on every rebuild; the build fails if any disagree. Interactive visualisation uses Phylocanvas^44^ for phylogenetic trees, Cytoscape.js^45^ (with *ELK* and fcose layouts) for pathway and network graphs, 3Dmol.js^46^ for protein structures from *RCSB PDB*, igv.js^47^ as the embedded genome browser, and MapLibre GL for maps^48^. The in-browser *BLAST* page runs a purpose-built Smith-Waterman kernel^49^ compiled to WebAssembly (Rust); no sequence data leaves the browser.

### Agentic Development

The processing pipeline as well as user interface was constructed primarily via agent-led development from iterative development and prompting by the lead author. Agentic development was executed primarily by the Pi coding agent^50^ with Ziphu AI *GLM*-5.3 and *GLM* 5.3-Flash models^51^, plus a locally hosted Qwen 3.6 with 27B parameters^52^. Generation parameters for local models followed the provider’s recommendations and were used via llama.cpp^53^ and sglang^54^ servers.

### Literature Analysis

To prevent copyrighted literature from being used for training, all literature analysis was performed using the local AI model Muse Glimmer^55^; this was run on SGLang with DFlash v2^56^ enabled, hosted on the University of Otago’s Aoraki Cluster located on the South Island of New Zealand, using 99.3% low-carbon energy, see https://www.otago.ac.nz/sustainability/doing/sdg/contribution#sdg-7-affordable-and-clean-energy. All literature, including scientific articles in *PDF*, media and government reports, is converted to Markdown format for processing. Literature is then classified as either a scientific journal article, preprint, thesis, news and media report, or other, by the *LLM* (Muse Glimmer).

The analysed corpus comprises 1,366 publications and reports. Literature analysis is performed using a purpose-built agentic multi-step approach: papers are registered with metadata (*DOI*, journal, year) and duplicate/junk guards; a per-paper analysis pass extracts genes, species and mutations with their roles; and follow-on passes extract quantified key findings, typed gene claims (e.g. *in vitro*, perturbation, sequence variation), and quote-grounded molecular mechanisms. Each stage is resume-safe (per-paper ledgers), and every extracted claim carries its source paper identifier and, for mechanism and spread items, a verbatim quote. Independent verification passes adjudicate extraction quality — quote-fidelity adjudication of extracted items currently passes 86% of items as faithful, and disputed items are flagged in the released data. Deterministic checks for additional mentions of genes and species are also performed using regular-expression scans with curated symbol-alias maps, complementing the LLM extraction; text-mention evidence is labelled separately from studied-role evidence and never presented as the stronger tier.

Extracted evidence is aggregated into a gene registry (4,277 host genes with per-gene paper counts, evidence classes and species links), per-gene and per-species synthesis pages, and the curated host gene panel. Species names are canonicalised against the site taxonomy with synonym folding; gene symbols against an alias map (e.g. *RIG-I*/*DDX58, MDA5*/*IFIH1*).

Questions and fact retrieval queries were generated via Pi development sessions depending on the types of data desired for the Roost website. All intermediate outputs are stored as per-paper *JSON* records and append-only ledgers (analyses, claims, mechanisms, verification verdicts), which serve as the source for every literature number shown on the site and reported here.

As an additional safety measure, mammalian mutations are censored from the website and data downloads.

### Detection Mining

To tabulate which species have had positive detections, we mine detection events from a variety of sources, updated periodically. These include *WOAH WAHIS*^11^, *USGS* WHISPers^17^, *FAO EMPRES*-i^18^, *USDA APHIS* wild-bird surveillance, the Canadian Wildlife Health Cooperative, UK *DEFRA*, Ireland *DAFM*, South Africa *OCIMS*^19^, ImmunomeDB^25^, Wildlife Health Australia^21^, and New Zealand *MPI* / Biosecurity NZ^20^. We found sources via web search, and Pi developed mining and extraction scripts that pull only new data to avoid excessive traffic spikes on remote hosts, and to deduplicate sightings and detection events while normalising fields (species names, dates, administrative geography, subtype/strain) across sources. The collated dataset currently holds 44,871 detection records covering 718 detected host species and more than 210 million individual animals, the large majority of which are poultry. Some detections are only found in news articles or government reports, such as New Zealand detections, and those are sourced from the literature analysis pipeline above. These are recorded and plotted on a built-in map, but Roost should only be a secondary source for up-to-date detections. The collated, deduplicated detection data is available for download.

### Genomes and Annotation

Genome assemblies and gene annotations were retrieved from *NCBI*^22,23^. We stage one assembly per species (3,894 assemblies, 2,093 non-redundant species; 689 chromosome-level assemblies), preferring the most recent representative assembly, and verify each download against a *SHA*-256 manifest; raw FASTAs are retained compressed. Predicted gene annotations are available for 302 assemblies covering 255 species. Additional protein sequences come from UniProt^24^, taxonomy and vernacular names from Birds of the World/Clements^26^ with name resolution via *GBIF*^27^, conservation status from the *IUCN* Red List as mirrored in Wikidata (property *P141*; 683 of 718 detected species assessed), and protein structures from *RCSB PDB*^28^. Curated facts from Avian Immunology^29^ are used as reference material only.

### Gene Panel and Presence Calls

The host gene panel (1,828 genes) is curated from the literature-derived candidates plus a UniProt/ GO immune-function backbone (GO:0002376), with legacy symbol aliases folded to canonical *HGNC* symbols. Each gene carries a curation importance class — *critical* (loss or gain directly tied to HPAI outcome: *DDX58*/*RIG-I, IFIH1*/*MDA5, ANP32A*), *high* (strong literature evidence for outcome relevance), or *standard* — and a curation tier (Supplemental Table 1).

Each panel gene is searched across every staged genome, producing one presence call per gene per genome at six evidence levels: E5, the gene is recorded in the *NCBI* annotation (search results are retained as corroboration); E4, no annotation but a sequence search finds the full gene; E3, a partial/fragment hit; E2, searched and not found; E1/E0, genome not searchable at that locus. Sequence search uses protein-level *DIAMOND*^57^ against reference orthologs (chicken and duck where available), supplemented by *NCBI BLAST*+^58^ blastn and tblastn, with *ORF*-coverage summarised from grouped hits via *EMBOSS* getorf; alias calls for the same gene and assembly are merged best-evidence-wins. The current matrix holds 2.95 million calls across 2,388 genomes; alias-folded per-genome detection rates are 83.5% (median 83%, *IQR* 78–89%) for Tier 1 host dependencies and 87.6% (median 89%) across all panel genes.

#### Orthology

To place the panel in a comparative frame, we ran OrthoFinder v3.1.5^59^ on a 463-species avian set drawn from the staged genomes, yielding 40,494 orthogroups. We mapped panel genes to orthogroups by primary assignment, with a chicken cross-check: 992 panel genes resolve to 721 orthogroups.

### Quality Control

Beyond the extraction-verification passes above, the build is guarded by deterministic cross-checks (21 artifact-vs-artifact and artifact-vs-page totals, including species, gene-registry, presence and download counts), a browser test suite covering interactive routes and accessibility (*WCAG* 2.1 AA scans of every canonical route), and a translation-integrity pipeline for the site’s seven languages. Manuscript numbers were audited paragraph by paragraph against the live artifacts; the audit trail and regeneration scripts ship with the repository.

## AI Statement

See methods.

## Data and software availability

Code is available at http://github.com/GenomicsAotearoa/Roost. Synthesized and public data are available on the website at https://josephguhlin.com/Roost/#/downloads. Original sources are available online or via publishers’ websites.

